# Reverse-engineering flow-cytometry gating strategies for phenotypic labelling and high-performance cell sorting

**DOI:** 10.1101/278796

**Authors:** Etienne Becht, Yannick Simoni, Elaine Coustan-Smith, Maximilien Evrard, Yang Cheng, Lai Guan Ng, Dario Campana, Evan Newell

## Abstract

**Motivation:** Recent flow and mass cytometers generate 1,000,000 single cell datasets of dimensions 20 to 40. Many tools facilitate the discovery of new cell populations associated with diseases or physiology. These discoveries require the identification of new gating strategies, but gating strategies become exponentially harder to optimize when dimensionality increases. To facilitate this step we developed Hypergate, an algorithm which given a cell population of interest identifies a gating strategy optimized for high yield and purity.

**Results:** Hypergate achieves higher yield and purity than human experts, Support Vector Machines and Random-Forests on public datasets. We use it to revisit some established gating strategies for the identification of Innate lymphoid cells, which identifies concise and efficient strategies that allow gating these cells with fewer parameters but higher yield and purity than the current standards. For phenotypic description, Hypergate’s outputs are consistent with fields’ knowledge and sparser than those from a competing method.

**Availability and Implementation:** Hypergate is implemented in R and available at http://github.com/ebecht/hypergate under an Open Source Initiative-compliant licence.

## Introduction

Analysis of low-dimensional cytometry data has historically relied on a procedure known as manual gating. Gating is the process of setting thresholds on measured parameters, to filter out unwanted cells until only a population of interest (PoI) is left. Cytometers can only measure a limited number of parameters (10-20 for modern flow cytometers, around 40 for mass-cytometry). It is thus crucial to carefully select the proteins that will be measured (a process known as ‘panel design’). Cytometry panels typically include a large number of well-characterized ‘lineage markers’ which help identifying cell populations. In addition, panels may include less-documented proteins, that may be relevant to certain fields or purely exploratory.

Gating in a low-dimensional setting is straightforward. It usually consists of dichotomizing up to each parameter, which divides the features space into up to 2^n^ volumes. Cell populations are defined by the events contained in each volume. Quantifying these populations across samples allow the identification of PoI that are associated with a study’s objectives. In this low-dimensional setting, reporting on the PoI is typically done by reporting the combination of expressed or absent markers that identify it (Figure 1A).

**Figure 1.**
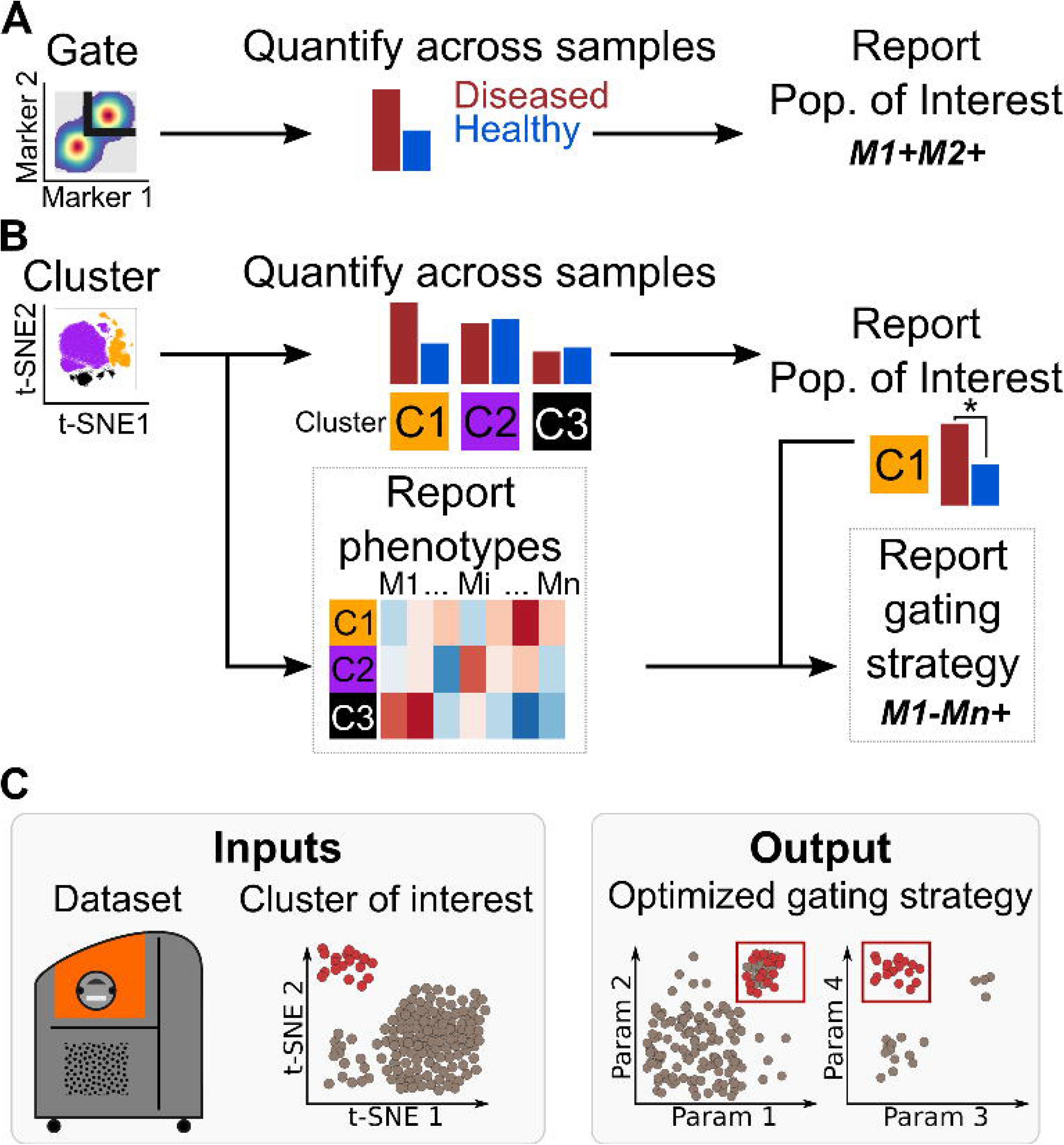
Schematic description of the analysis of **A**) a low-dimensional or **B**) high-dimensional cytometry dataset. Framed are steps for which only limited software assistance exist. **C**) Summary of Hypergate’s workflow. The red-colored events represent a ‘population of interest’.

Classical gating becomes complex in a high-dimensional setting, as the number of volumes created by exhaustively dichotomizing parameters quickly outgrows the typical number of single cells profiled by these methods (Cheng et al. 2016). Exploratory analyses therefore rely on tools designed to operate directly on the high-dimensional data. These tools include dimensionality reduction techniques for visualization (reviewed in Saeys et al. (2016)), clustering techniques to define cell populations in an unsupervised manner (benchmarked in Weber et al. (2016)), trajectory detection methods that output an ordering of the high-dimensional data (reviewed in Cannoodt et al. (2016)) and more recently supervised methods that identify cell populations whose frequencies differ across sample types (Lin et al. 2015; Arvaniti et al. 2017; Lun et al. 2017). While these tools help identifying PoI in a high-dimensional setting it remains necessary to describe their phenotype in a way that resonates with current knowledge and a protocol to identify them in independent experiments (Figure 1B).

The complexities of both tasks augment with dimensionality and software to facilitate them is needed. Phenotyping a cell cluster among *N* cell clusters in a *P* dimensional space will involve (*N*-1)\**P* comparisons. For typical mass-cytometry values of *P* = 40 dimensions and *N* = 10 clusters, up to 360 comparisons are required to annotate one cluster alone. Manually annotating cell phenotypes in high-dimension is thus a time consuming process. Finding an optimal gating strategy is theoretically even harder, as the search space grows exponentially with dimensionality. We for instance show in the *Methods* that the search space of possible rectangle-shaped gating strategies (after filtering out trivially non-optimal ones) for a small PoI of size 100 in a 40-dimensional dataset is higher than 10^146^. Since finding a gating strategy also provides a phenotypic label, we focused on developing a method to identify gating strategies when given a PoI.

## METHODS

### Datasets

For benchmarking purposes, we downloaded public datasets with manual gating annotations that were previously collated by Weber and Robinson (Weber et al. 2016) from FlowRepository (Spidlen et al. 2012) accession number FR-FCM-ZZPH. Briefly, we used three high-dimensional flow cytometry and four mass-cytometry datasets. One flow cytometry dataset (‘FlowCAP_ND’) originates from the FlowCAP-I challenge (Aghaeepour et al. 2013) and profiles the blood of 30 healthy donors. The other two profile either the blood (Mosmann et al. 2014) or the bone marrow (Rundberg Nilsson et al. 2013) of one healthy donor. The four mass-cytometry datasets consist of two human bone marrow datasets (Levine et al. 2015) and two mouse bone marrow datasets (Samusik et al. 2016). We also analyzed two cord-blood samples enriched for human innate lymphoid cells (ILCs) available from FlowRepository (FR-FCM-ZYZX, samples Cord Blood #1 and #2). From the ILC dataset we pre-gated on live (Cisplatin^-^) singlet (DNA^int^) immune (CD45^+^) cells prior to analysis.

### Objective function

Hypergate optimizes for the Fβ score, where Fβ = (1+β^2^)(purity. yield)/(β^2^.purity + yield). It can be reformulated in terms of the number of True Positive (TP), False Negative (FN) and False Positive (FP) events as Fβ=(1+β^2^). TP / ((1+β^2^)TP + β^2^.FN + FP). Fβ is in general bounded by 0 (yield or purity is 0) and 1 (yield and purity are both 100%). Otherwise it is bounded by min(purity,yield) and max(purity,yield). It is also known at the harmonic mean of purity and yield. Unlike the arithmetic mean it lies closer to the lowest of the minimum of purity and yield rather than at their barycentre.

### Hypergate operation (Supplementary Figure 1)

### Overview

At initialization, the gating strategy is an infinite rectangle in every direction for every parameter (the whole feature-space is thus initially gated-in). Yield is thus 100% and purity N+/N, where N is the total number of events in the dataset and N+ the number of events of interest. From there, Hypergate uses two basic operations. We refer to the first one as a contraction, where a given threshold is made more stringent (thus shrinking the hyperrectangle on one of its faces). A contraction will decrease the number of cells gated in, which may increase purity at the cost of yield. We refer to the second move as an expansion, where the effect of enlarging the hyperrectangle so that it contains a gated-out event of interest is evaluated. Expansions increase the size of the hyperrectangle, and thus may increase yield at the cost of purity. Hypergate performs a succession of these basic operations to gradually increase Fβ until no move is able to locally increase it further, at which points it terminates.

### Parameters

Hypergate takes as input an expression matrix *X* of *N* rows (events) and *P* columns (parameters), a boolean vector *S* of size *N* designating the subset of interest (or the subset of “positive” events) and *β* which specifies the yield versus purity weighting (default to 1, i.e. equal weighting).

Internally, the algorithm keeps track of both the lower and upper cut-offs on each parameter (two numeric vectors C^-^_1≤j≤P_ and C^+^_1≤j≤P_), and the Boolean state *B*_*1≤i≤N*_ which specifies for each event whether it is currently gated-in or gated-out.

### Initialization

At initialization, the lower (respectively upper) cut-off of every (parameter, direction) pair *p* is set at min_1≤i≤N_(pi) (resp. max_1≤i≤N_(pi)) and *p* is set to inactive.

### Updating the state vector

For each datum x_i_ = (y_i,1_,…, y_i,j_,…, y_i,p_), the corresponding state B_i_ is set to True (or “gated-in”) if and only if ∀j | 1≤j≤P, C_j_^-^≤y_i,j_≤C_j_^+^, and to False (or “gated-out”) otherwise. This corresponds to points within the hyperrectangle, including the boundary.

### Contractions

A contraction is the shrinkage of the hyperrectangular gate, which results in less events being gated-in. In order to evaluate only a meaningful and computationally-manageable subset of all the possible contractions, we use the following three ideas: i. discretization: any two shrinkages on a same side of the hypperctangle (i.e. a single (parameter, direction) pair) that both lie between the same two consecutive events will be equivalent in terms of the gating state *B* it produces. We thus only evaluate threshold values that are present in the dataset. ii. An hyperrectangle that would have no gated-out event of interest on one of its faces (including their boundaries and after discretization) would lead to a suboptimal gating, as the closest more stringent threshold corresponding to the expression of an event of interest on this channel would lead to the same yield but increased purity and thus increased Fβ. We thus only evaluate contractions on each parameter for values expressed by an event of interest. iii. In order to avoid combinatorial explosion, we only evaluate one-dimensional contractions. In summary, we evaluate one-dimensional contractions whose thresholds correspond to gated-in events of interest.

### Expansions

An expansion increases the size of the hyperrectangular gate, which results in more events being gated-in. As for contractions, the expansion(s) that maximize Fβ given a current stage of the hyperrectangle and after discretization have events of interest on their faces. For a gated-out event of interest x = (y_1_,…, y_j_,…, y_p_) and a given hyperrectangle H=(C_1_^-^,…,C _p_^-^, C_1_^+^,…,C _p_ ^+^), the inclusion of x_i_ into the gate will lead to an updated hyperrectangular gate H’=(min(C_1_^-^,y_1_),…, min(C _p_ ^-^,y _p_), max(C _1_ ^+^,y_1_),…, max(C_p_ ^+^,y_p_)). Given H’, we compute a new state B’, associated with a new score Fβ’. The gated-out event of interest associated with the highest-scoring expansion (if any) is used to update the hyperrectangular gate.

### Operations’ order of priority

Hypergate prioritizes expansions over contractions of an active (parameter, direction) pair over contractions of an inactive (parameter, direction) pair. This is equivalent to first maximizing yield as long as Fβ increases, then maximizing purity, and as a last resort using a new channel in the gating strategy.

### Termination

The algorithm terminates when no move increases Fβ or if the last channel added did not contribute more than a user-specified threshold (default 0).

### Time complexity for the brute force approach

The design of Hypergate is justified by the following considerations which show that a brute force evaluation of hyperrectangles is impractical: for a given parameter P, the hyperrectangle is defined by a lower threshold C _p_ ^-^ and an upper threshold C _p_ ^+^ with C_p_ ^-^≤ C _p_+. If we restrict the possible thresholds to values expressed by one of the N^+^ events of interest (as described in the ***Contractions*** section), then there are N^+^(N^+^-1)/2 = N^+^ choose 2 possible pairs for (C _p_ ^-^ C _p_ ^+^), provided the N^+^ events have distinct expression values for the parameter P. For a given dataset of p parameters with N^+^ events of interest, the number of hyperrectangle to evaluate by brute force is in the order (as here we are not accounting for possible duplicated values) of (N^+^(N^+^-1)/2)^p^ For a small population of 100 events and 40 markers, the search space is for instance higher than 10^146^ Hypergate proposes an approximate solution for this problem.

### Ranking of output’s parameters

Given a cytometry dataset and a subset of interest, Hypergate outputs an Fβ score ranging from 0 to 1 using a finite number of (parameters, direction) pairs. In order to evaluate which ones are the most significant within the output, we evaluate the difference between the Fβ score at the end of the optimization procedure, and the one obtained when the same gating strategy is used but omitting the use of one parameter. Omitting a parameter will necessarily lead to a decrease in Fβ, and the difference δFβ is used as a proxy for parameter importance. This metric could fail to capture certain effects, for instance if two parameters are redundant (so that removing one will have little effect and thus both will appear relatively unimportant, yet removing both will not). It nonetheless appears to perform well in practice.

### Transformations

For the public manually-gated datasets, we used the data as transformed by Weber et al. (namely x→ arcsinh(x/5) and x→arcsinh(x/150) for mass and flow cytometry datasets, respectively). For the ILC dataset we used the transformation x→arcsinh(x).

### t-SNE

For the ‘Samusik_01’ dataset, we randomly sampled up to 100,000 events, while retaining the manually-gated populations frequencies. We then used the Rtsne function from the Rtsne R package, which implements Barnes-Hut t-SNE (van der Maaten et al. 2008; Van Der Maaten 2014), using up to 1000 iterations, and a perplexity of 30. For the ALL (respectively ILC samples), we ran t-SNE independently on each sample, using pre-gated CD19^+^ cells (respectively DNA^int^ CD45^+^Cisplatin^-^ cells) without down-sampling using up to 1000 iterations, and a perplexity of 30.

### Reproducing manual gating strategies (Figure 1B and Figure 3)

We referred to the Supplementary Figure 5 of the Samusik et al publication (Samusik et al. 2016) to identify the channels that were used by the authors to gate on each cell population. We formalized this graphical depiction of the gating strategy into a tree using mathematical logic (a population positive for two markers A and B will be noted as (A+ *and* B+), and the rest will be noted (A- *or* B-)). In figure 1B, we used the convex hull of gated-out events on each of the sequential bi-axial plots to reproduce the gating steps.

**Figure 2.**
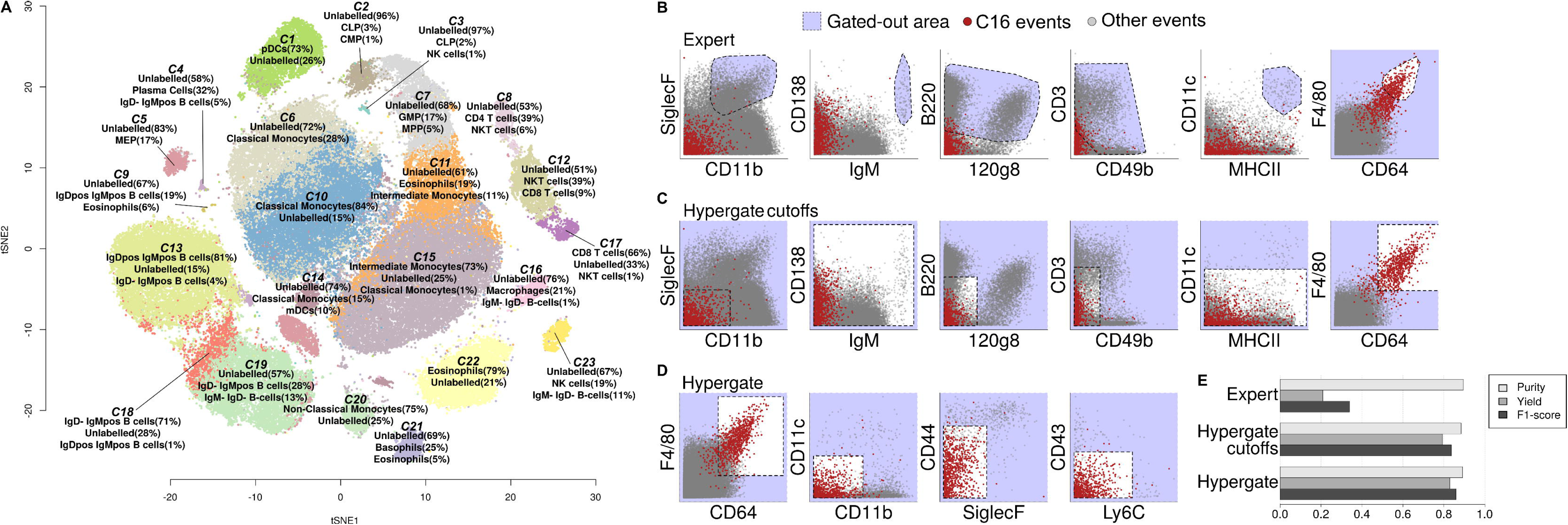
**A**) t-SNE representation of the ‘Samusik_01’ dataset color-coded by cluster identity output by Phenograph. **B**) Reproduction of the gating strategy defining Macrophages according to Samusik et al, using cut-off values defined by the authors. The blue area is excluded at each step and the non-excluded events are represented on the next step. The red events correspond to t-SNE-defined macrophages. **C**) Reproduction of the gating strategy defining Macrophages according to Samusik et al, using cut-off values computed by Hypergate. **D**) 8-channels gating strategy for t-SNE-defined Macrophages identified by Hypergate. **E**) Comparison of the three gating strategies in terms of purity (light gray), yield (dark gray) and their harmonic mean (F1-score, black).

### Supplementary Video

We chose to illustrate Hypergate’s operation on the “Non classical monocytes” population of the Samusik_01 dataset as there is a high concordance between this population and the corresponding t-SNE cluster. We trained Hypergate on the whole dataset using this entire cell population as the population of interest, with a beta parameter of 1 (default value).

### Hypergate applied to t-SNE-like macrophages

We applied Hypergate using the authors’ gating strategy on the cluster of macrophages defined by t-SNE (as highlighted in Fig 2A), using a value of 1 for the *beta* argument (default value), and using either the parameters chosen by the authors or letting Hypergate identify the parameters to use. In the latter case, we restricted the gating strategy to the 8 most significant parameters (see ***Ranking of output’s parameters***).

### Automated phenotypic description of cell clusters

To obtain cell clusters, Phenograph was ran on the ‘Samusik_01’ dataset, using a k=30 nearest neighbours (default parameter). This resulted in 23 clusters. From each cluster, we sampled up to 1000 positive events, the corresponding proportion of negative events, and ran Hypergate on the resulting data subset. We then scored the importance of the parameters’ in Hypergate outputs for each cluster (see ***Ranking of output’s parameters***), and labelled each cluster with its four most defining parameters.

### Automated phenotypic description of manually-gated cell populations

From each cell population, we sampled up to 1000 positive events, the corresponding proportion of negative events, and ran Hypergate on the resulting data subset. For each Hypergate output, we computed each parameter’s significance (see ***Ranking of output’s parameters***). We then used the *wordcloud* R package (version 2.5) using the square-root of the parameters’ significance to map the size of the words, with a four-to-one ratio between the maximum and minimum size.

### MEM scores

MEM scores were computed according to the formula published by Diggins et al. (2017) using a de-novo R script.

### Binary classification

From each dataset and for each cell population manually gated by the authors’, we sampled 1000 events if the population was larger than 2000 events, or half of the events otherwise. We also sampled a corresponding proportion of negative events. We then trained 6 classifiers on this training set: Hypergate, Nelder-Mead optimization of an hyperrectangle, four Support Vector Machines, and a Random Forest. For Nelder-Mead optimization, we used the *optim* function of the *stats* R package (version 3.4.2). For Random Forests, we used the *randomForest* function of the *randomForest* R package (version 4.6-12) with default parameters. For Support Vector Machines, we used the *svm* function of the *e1071* R package (version 1.6-8), using default arguments except for the ‘kernel’ argument (which we set either to ‘linear’ or ‘radial’) and the ‘class.weights’ argument which we set either to 1 (constant) or to the inverse frequency of the classes (thus weighting the rarer class more). We then used these 6 models to predict the class of the left-out data (the ‘test set’), and computed the corresponding F1-scores and accuracies. This workflow was repeated for each of the 95 populations defined by the authors of the datasets used. We used the same algorithms to analyze the ALL data.

### Acute Lymphoblastic Leukemia samples

### Procurement and processing of samples

Bone marrow samples from 5 patients with newly diagnosed acute lymphoblastic leukaemia (ALL), aged 1-23 (median, 7) years, were obtained at diagnosis and during treatment. Bone marrow samples obtained during treatment from another 5 patients, aged 1-12 (median, 5) years, that were leukemia-free by both flow cytometry and polymerase chain reaction-amplification of antigen receptor genes, were used as a reference. Samples were obtained following informed consent and approval from the Domain Specific Ethics Board governing Singapore’s National University Hospital. Mononucleated cells were labelled with fluorochrome-conjugated antibodies as previously described (Coustan-Smith et al. 2011).

### Identification of blasts at diagnosis

We examined the expression of ALL markers color-coded on t-SNE biplots at the diagnosis time point to manually identify the malignant population. This information was validated by an expert haematologist.

### Hypergate training and translation to follow-up samples

Each diagnosis sample was concatenated with data from 5 leukaemia-free bone marrow samples. We then trained Hypergate to obtain a gating strategy that identifies the malignant population, using a beta parameter of 1 (default value). We applied these gating strategies to follow-up samples using the same cut-offs (as these clinical samples present little technical variability).

### Statistical tests

We used paired t-tests assuming unequal variances to compare F1-scores and accuracies across cell populations for each binary classifier against Hypergate. We used a test against the t-distribution to assess non-null correlation between log-frequencies measured by Hypergate and other classifiers and an expert haematologist.

## Results

### Development of the Hypergate algorithm

We developed Hypergate (for automated HYPERrectangular GATE), an algorithm which given a PoI outputs a corresponding gating strategy (Figure 1C). Hypergate operates by finding a hyperrectangle (or high-dimensional rectangle) that specifically encapsulates the cell cluster of interest. It does so by iteratively modifying the boundaries of the hyperrectangle while optimizing for an Fβ-score (by default the F1-score, the harmonic mean of yield and purity). Hypergate is a deterministic algorithm and thus enables easily replicable results. Its operation is depicted in Supplementary Figure 1, detailed in the Methods, and an example of its execution is shown in Supplementary Video 1.

### Hypergate overcomes issues associated with manual gating strategies

Figure 1A represents a publicly-available (Samusik et al. 2016; Weber et al. 2016) mass-cytometry mouse bone-marrow dataset manually-annotated for 24 haematopoietic populations. We analyzed it using t-SNE (van der Maaten et al. 2008) for dimensionality reduction and Phenograph (Levine et al. 2015) for clustering. Phenograph identified 23 clusters. For each cluster, we overlaid on the t-SNE space the 3 most frequent manually-gated cell types. This analysis highlights various pitfalls associated with manual gating and that are particularly manifest in high-dimension. For instance, the number of events belonging to some cell populations seems underestimated by a factor of 3-4 by manual gating as compared with well-delineated Phenograph clusters (clusters C2 enriched in Common Lymphoid Progenitors and C16 enriched in Macrophages). In addition, examination of cluster C23 highlights that NK cells are likely contaminating the manual IgM^-^IgD^-^ B cells, and examination of C11 suggests that Eosinophils contaminate the Intermediate Monocytes gate. Contaminations can be especially troublesome for the interpretation of data from FACS-sorted cell populations, such as transcriptomics. Nonetheless, some clustes are highly concordant with manual labels (C1 with plastmacytoid Dendritic Cells, C20 with Non-Classical Monocytes). Projection of manual-gating labels on the t-SNE map is shown in Supplementary Figure 2 and highlights the same limitations.

Projection of the macrophage-rich cluster C16 on the gating strategy used by the authors to identify macrophages showed that a low CD3^-^ threshold and too stringent F4/80^+^ and CD64^+^ thresholds contributed most to this underestimation (Figure 2B). Applying Hypergate this C16 cluster produced a wider CD64^+^F4/80^+^ gate that accommodated the large background expression of these markers (Figure 2C). Interestingly, 3 out of 12 markers used by the authors to define macrophages were not contributing to the final output (CD138, IgM, and MHCII). We thus ran Hypergate in an unconstrained mode, where parameters’ selection solely depends on the algorithm. This led to a shorter 8-parameters strategy (Figure 2D) that enabled comparable purity, but higher yield (Figure 2E, 20.9% for the authors’ gating versus 79.3% and 82.9% for supervised and unsupervised Hypergate respectively) and the highest F1 score (0.334 versus 0.836 and 0.859 respectively. Interestingly, while crucial to the gating strategy, CD64 thresholding was more loosely enforced in the final strategy, primarily due to the use of CD44 (Supplementary Figure 3) which appears as a useful exclusion marker to gate on macrophages in this context.

**Figure 3.**
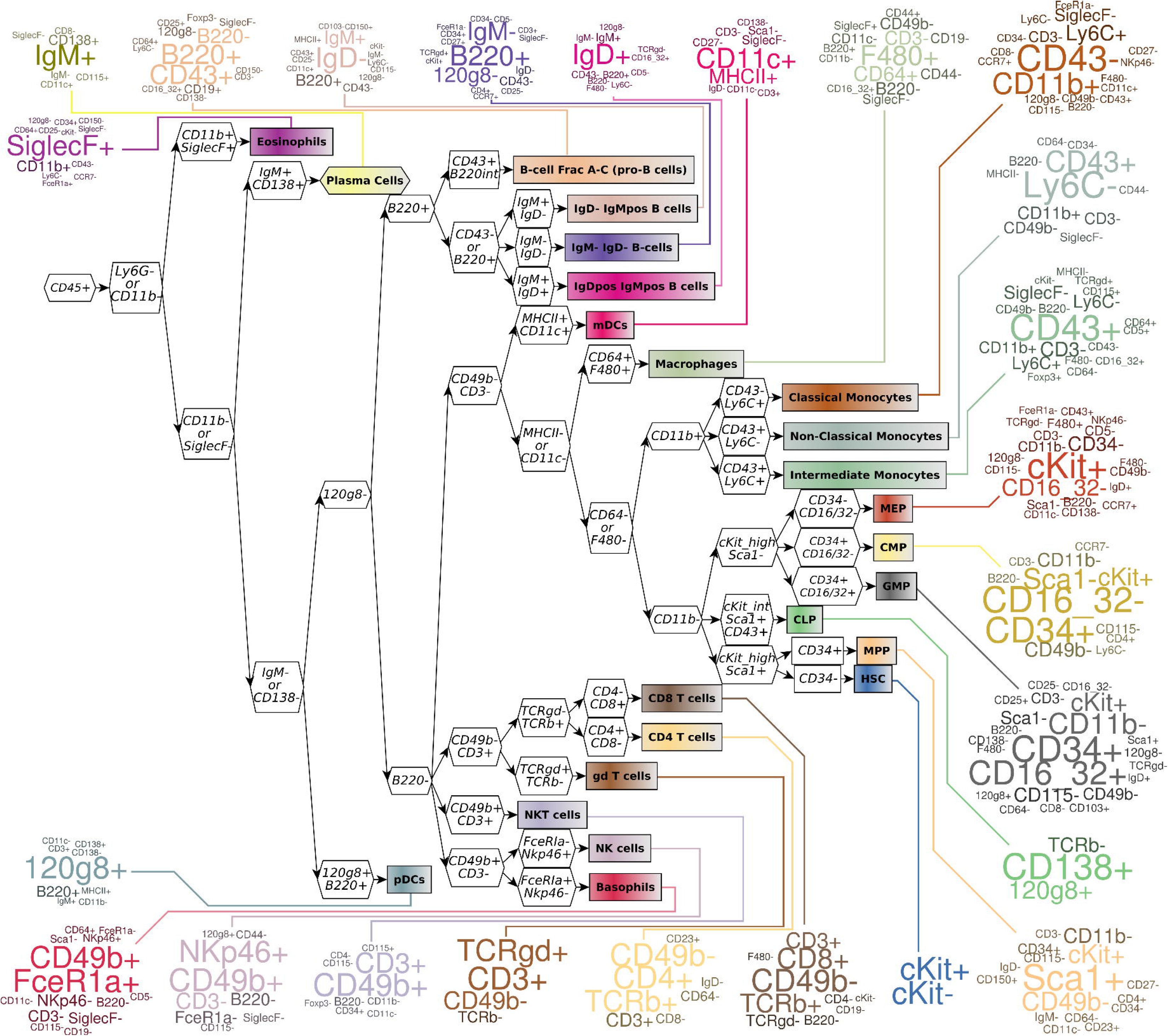
Comparison of Hypergate-computed cell phenotypes and Samusik et al. gating strategies. The middle part of the graph (the gating tree) summarizes the authors’ gating scheme. The leaves of the gating tree represent populations. Wordclouds around the populations show the parameters selected by Hypergate to gate on each expert-defined population. Significance of each parameter is encoded by its size (bigger denotes more significant).

### Generation of parsimonious phenotypic descriptions

The fact that 5 out of the 8 markers identified by Hypergate to define macrophages-like cells are consistent with the experts’ selection suggested that our method could be used to output concise phenotypic descriptions of cell populations, by measuring the relative contribution of each marker to the computed gating strategy (see Methods). The phenotype of C16 macrophages-like cells would for instance read as F4/80^+^CD64^+^CD11b^-^CD11c^-^ SiglecF^-^CD44^-^Ly6C^-^CD43^-^. An informed interpretation of this gating strategy would be to enrich for macrophages (F4/80^+^CD64^+^) and to exclude monocytes (using CD11b^-^CD43^-^Ly6C^-^), dendritic cells (CD11c^-^) and eosinophils (SiglecF^-^) and non-macrophage myeloid cells (CD44^-^). It shows that excluding T cells (using CD3^+^) or B cells (B220^+^) is not necessary, as F4/80^+^CD64^+^ double positivity is sufficient to exclude virtually all of these cells (Supplementary Figure 4).

**Figure 4.**
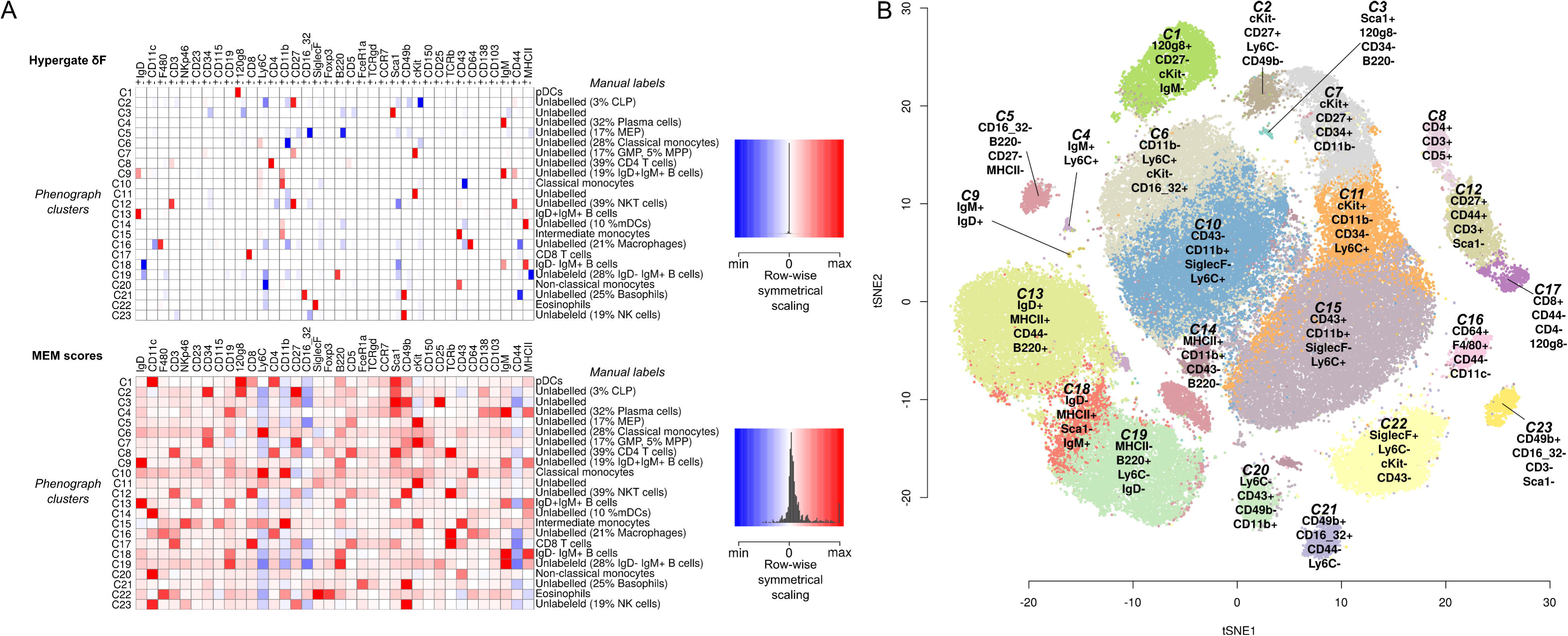
**A**) Heatmap showing phenotypic labels as computed by Hypergate (top) or MEM scores (bottom). Right panels feature histograms representing the densities of the corresponding scores. **B**) t-SNE map color coded by Phenograph clusters and annotated using Hypergate.

To study the accuracy of Hypergate’s phenotypic characterizations, we applied it systematically to each of the 24 populations defined by Samusik et al., and found that they were consistent with authors’ definitions (Figure 3). For instance, myeloid dendritic cells were algorithmically-labelled as CD11c^+^MHCII^+^, basophils as FceR1a^+^CD49b^+^CD3^-^SiglecF^-^NKp46^-^ and gamma-delta T cells as CD3^+^TCRgd^+^CD49b^-^TCRb^-^. Discrepancies were nonetheless found in labels for Hematopoietic Stem Cells and Common Lymphoid Progenitors due to low cell counts (3 and 59 events in this dataset, respectively). The MEM scoring method has recently been proposed as a metric weighting markers’ relevance to a cell cluster phenotype (Diggins et al. 2017). Unlike MEM whose output is of the same dimensionality as the input, Hypergate only selects for a subset of the parameters to characterize a cell population and thus produces more concise phenotypic characterizations which is important for their readability (Figure 4A). Moreover, Hypergate can identify markers as intermediately expressed (e.g. pro B-cells are accurately as expressing intermediate levels of B220, Fig 3). Combining Hypergate, t-SNE and Phenograph clustering allowed for an unsupervised pipeline that output cell clusters annotated with human-readable labels (Figure 4B). Cluster C8 was for instance labelled as CD4^+^CD3^+^CD5^+^F4/80^-^ (CD4 T cells), various B cell clusters were defined by combinatorial expression of IgD and IgM, and C21 as CD49b^+^CD16_32^+^CD44^-^Ly6C^-^ (Basophils).

### Performances as a binary classifier

To evaluate the performance of our method in terms of producing high efficiency gating strategies, we considered it as a binary classifier. We benchmarked it on 95 cell populations defined across 7 public datasets (Weber et al. 2016). For each population, we trained Hypergate, a general optimization method to solve the same optimization problem (Nelder et al. 1965), Support Vector Machines (SVMs) and Random Forests (RFs) on a subset of the data and assessed each classifier’s performance on the remaining unseen data. Hypergate led to higher F1 values than other method (Figure 5A), and higher total accuracy than SVMs and Nelder-Mead optimization, comparable to RFs despite optimizing for the F1 score (Figure 5B). These results suggest that Hypergate is able to output high-performance gating strategies. Furthermore, unlike Nelder-Mead optimization, SVMs and RFs, Hypergate’s outputs are straightforward to interpret in terms of each cell population’s phenotype.

**Figure 5.**
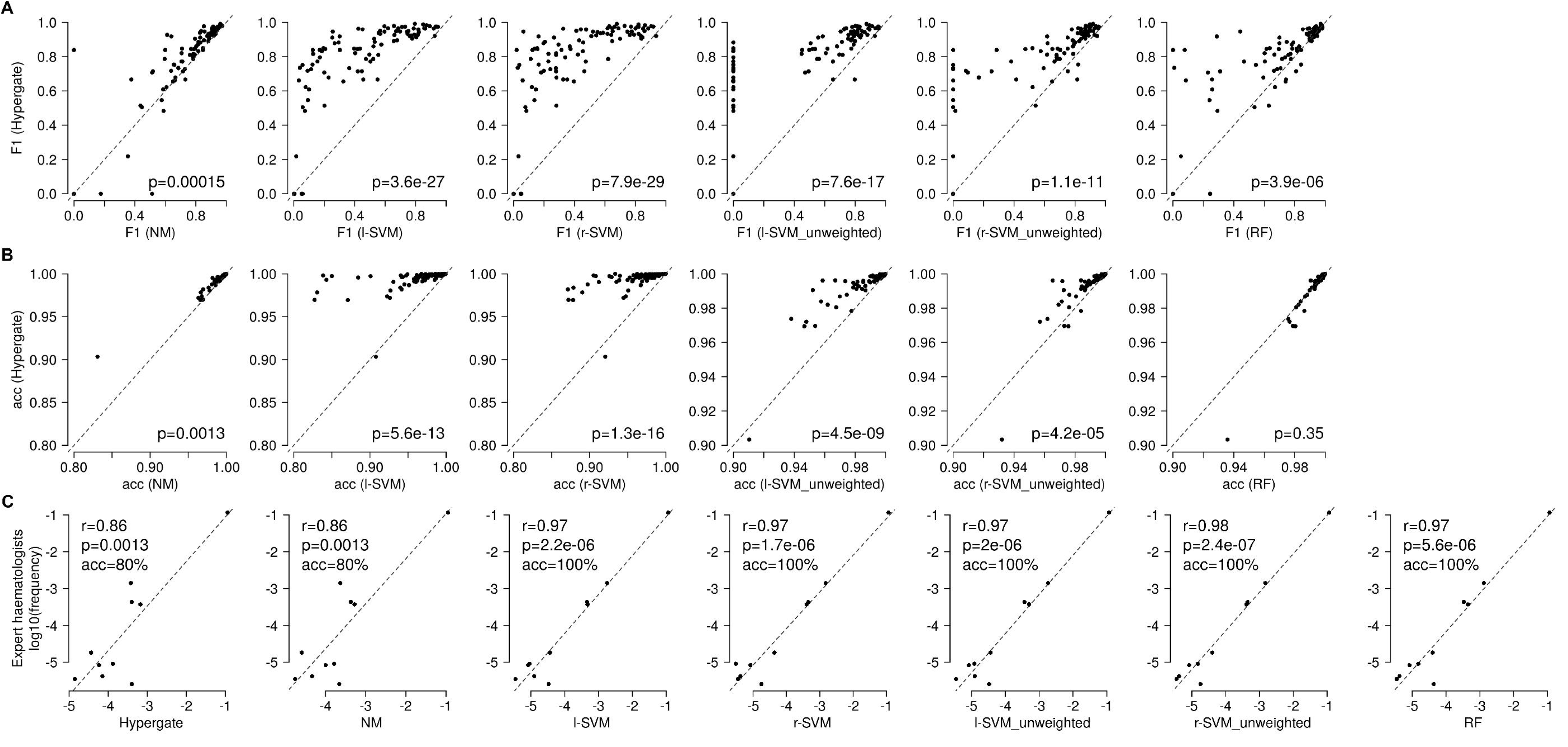
Biplots comparing the **A)** F1 scores and **B)** accuracy in binary classification across 95 populations and 7 datasets for Hypergate, Nelder-Mead optimization, four support vector machines and random-forests. The dashed-line represents the first diagonal (line of slope 1 that crosses the origin). p-values are computed from a paired Student t-test assuming unequal variances. **C)** log10-proportion of malignant cells among PBMCs estimated either by expert haematologists (y-axis) or other classifiers (x-axis). Pearson correlation coefficients and the corresponding p-values are reported, as well as diagnostic accuracy (using expert diagnosis as a gold standard).

Binary classification has applications in clinical haematology. One of them is the diagnosis of Minimal Residual Disease (MRD) in the context of Acute Lymphoblastic Leukaemkia (ALL) (Coustan-Smith et al. 2011). MRD is evaluated after chemotherapeutic treatment and based on the quantification of remaining malignant cells in the bone marrow. Positive MRD (high frequency of surviving blasts) is associated with faster relapse. By using a combination of Hypergate and t-SNE, we reproduced this clinical workflow. We trained Hypergate to gate on malignant cells at diagnosis for 5 patients and applied them to follow-up samples (from 1 to 3 samples per patient, Supplementary Figure 5). The computed log-frequencies of malignant cells among total PBMCs significantly correlated with those measured by expert haematologists (Figure 5C, r=0.86, p=0.0013). Across all follow-up samples and using the usual 1/10000 malignant cell per PBMC for a positive MRD detection, we obtained 80% accuracy in diagnosing MRD, significantly higher than random chance (p=0.011). Although Hypergate has the advantage to define the malignant cells’ phenotype, SVMs and RF yielded higher linear correlation coefficients (r>0.96) and higher diagnostic accuracies (100%) across these 10 test cases. This is likely due to the ability of both methods to leverage the high-dimensional data for classification, which may be helpful in a setting where some biological variability between the training and test samples can be expected.

### Redefining a gating strategy for the isolation of cord-blood innate lymphoid cells

We challenged Hypergate on a practical immunology task: the gating of innate lymphoid cells (ILCs). ILCs are a recently-described lymphoid population that, akin to NK cells, do not rely on rearranged antigen receptors for their activation. ILCs are, like T helper (T_h_) cells, non-cytotoxic and cytokine-producing cells. The exact number of distinct ILC subsets is still debated, but the ILC2 (Th2-like) and ILC3 (Th17-like) subsets are consensual (Hazenberg et al. 2014; Eberl et al. 2015; Simoni et al. 2017).

On a cord blood sample first depleted for T and B cells using magnetic cell sorting and then profiled using mass-cytometry, t-SNE identified two well-delineated clusters corresponding to ILC2s and ILC3s (Figure 6A). The standard gating strategy for the identification of ILC2s or ILC3s requires more than eight surface markers (Figure 6B). ILCs are commonly defined as CD45^+^CD14^−^CD34^−^CD5^−^Lineage(FceR1,CD19,CD123)^−^CD94^−^ CD127^+^ and subdivided into CRTH2^+^ ILC2s and CRTH2^−^c-Kit^+^NKp44^+/–^ ILC3s (Figure 6B). Taking the corresponding t-SNE clusters as gold standards, these manual gating strategies resulted in F1 scores of 0.87 and 0.70, respectively.

**Figure 6.**
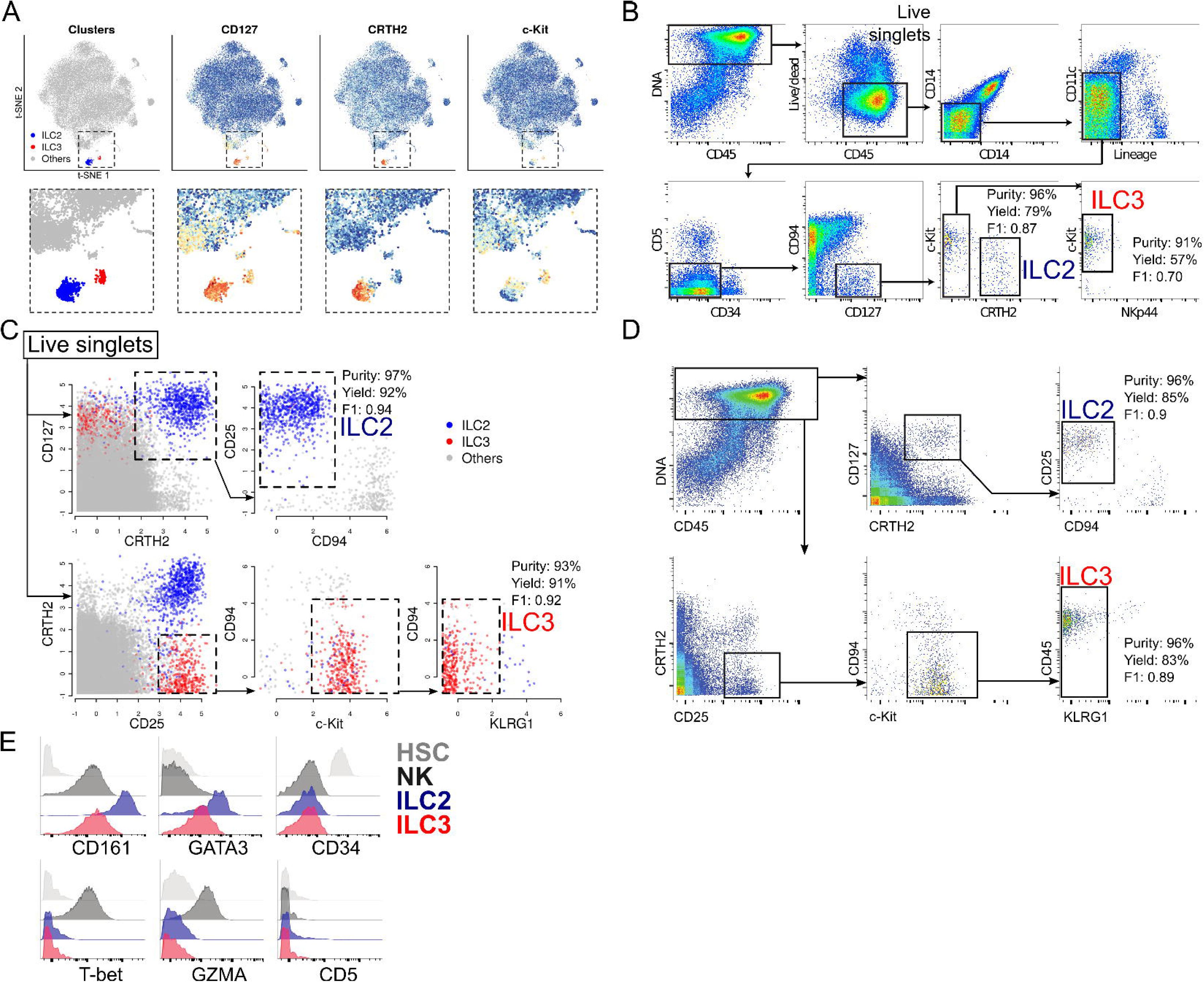
**A**) Identification of ILC2s ans ILC3s using t-SNE. **B**) Manual gating strategy for the identification of ILC2s and ILC3s. Purity, yield and the F1 score are defined using the t-SNE cluster as the ground truth. **C**) Gating strategies identified by Hypergate for ILC2s (top) and ILC3s (bottom). **D**) Blinded reproduction of the learnt gating strategy on an independent sample. **E**) Phenotype of the blindly-gated ILC subsets as well as NK cells and Hematopoietic Stem Cells as controls.

Hypergate identified shorter gating strategies for ILC2s and ILC3s (Fig 6C), both relying on CD94^-^ (exclusion of NK cells) and CD25^+^. In addition, ILC2s were identified as CD127**^+^** and CRTH2**^+^**, ILC3s as CRTH2^-^, cKit^+^ and KLRG1^-^ (Figure 6C). These strategies strikingly did not rely on the lineage ‘dump’ channel (FceR1/CD19/CD123 all conjugated to a Gd156 tag), nor Hematopoietic Stem Cells (CD34), monocytic (CD11c) or T cells (CD5) exclusion markers. Consistent with previous reports showing that not all ILC3-like cells express CD161 (Simoni et al. 2017; Li et al. 2018), this marker was not included in this Hypergate gating strategy. Both strategies resulted in F1 scores higher than 0.9.

A manual blinded replication of this gating strategy on an independent sample (Figure 6D) resulted and using de-novo t-SNE clusters as a gold standard resulted in a good purity and recovery of both subsets (F1 of 0.90 and 0.89 respectively for ILC2s and ILC3s). The gated-in cells featured expected levels of expression for various markers (CD34^-^CD5^-^T-bet^-^GZMA^-^), CD161^hi^GATA3^hi^ for ILC2s and CD161^int^GATA3^int^ for ILC3s, confirming their cellular identities (Figure 6E). These results demonstrate the ability of Hypergate to produce short, reproducible and high-performance gating strategies.

### Discussion and future works

Herein we introduce Hypergate as a new tool that extends current high dimensional cytometry analysis pipelines, allowing automated phenotypic annotation of cell subsets as well as the optimization of sorting strategies. Unlike manual gating, this approach is less prone to certain biases such as too conservative gating which leads to an underestimation of the frequencies, and is not biased against poorly documented markers nor in favour of well-documented ones.

Phenotypically labelling a cell subset either involves finding textbook labels matching its phenotype, or the identification of useful metrics to accurately illustrate the significance of each parameter in relation to the cell subset. The later case is notably suitable for new subsets and for subsets whose definitions are still controversial. Unlike other methods such as median or mean summarization or the MEM scoring (Diggins et al. 2017) approach, Hypergate outputs coefficients that are mostly null or close to null. These parsimonious phenotypic descriptions are faster to interpret which is relevant when the number of clusters is important (as it is commonly the case when studying immunological datasets).

Gating strategies are also widely used in cell sorting experiments, where a particular population is isolated for further studies. Sorting requires gating strategies that are short (as even modern cell sorters can only incorporate up to a dozen markers) and efficient. Purity is often the major criteria to avoid contamination of the population of interest, but it is hard to control for when using only a limited number of markers. Yield is harder to control for when using traditional manual gating, as the proof-reading process involved (known as “backgating”) focuses on improving purity rather than yield due to its non-commutativity. By identifying unbiased gating strategies established on large flow or mass-cytometry panels, Hypergate enables the task-specific definition of high-purity and high-yield gating strategies for sorting experiments. Applying our method to ILC subsets allowed us to identify them manually with high accuracy.

The current Hypergate implementation can only gate on a single cell cluster at a time. Gating strategies to identify many cell subsets could be implemented by optimizing for the mean F1 score across all subsets instead of a single measure, but the issue of matching subspaces to clusters arises. Weber et al. (2016) used the Hungarian algorithm for cluster-population assignment in their clustering benchmark, which could help automate this step while keeping the clusters distinct in the gating scheme.

Many studies (Aghaeepour et al. 2013; Samusik et al. 2016; Li et al. 2017) have proposed automated gating procedures whose focus is on the result of the gating procedure (the cell labels). Thus, while these methods can be powerful clustering tools, they do not easily allow mapping identified populations on independent data. We thus suggest that the method we describe herein will purposefully complement other available tools and could also prove useful to other types of high-dimensional data.

### Software availability

Hypergate is available for download at http://github.com/ebecht/hypergate

## Acknowledgements

We thank members of the Singapore Immunology Network and notably members of the E.N. laboratory. Shamin Li, Michael Fehlings, Melissa Chng, Michael Wong and Harsimran Singh are deeply acknowledged for their inspiring and insightful feedbacks. We would also like to thank those who produce, host and disseminate public data, which have been instrumental during the development of this work. Human donors who opted in to take part in research projects are thanked. This study was funded by A-STAR/SIgN core funding and A-STAR/SIgN immunomonitoring platform funding.

## Author contributions

EB and EN designed the study. EN supervised the study. EB developed and applied the algorithm. YS, ECS, ME, YC, LGN, DC and EN provided data. EB, YS, ECS, YC, DC analyzed the data. Each author contributed to writing the manuscript.

## Abbreviations

FACS: Fluorescence-activated cell-sorting
ILC: Innate Lymphoid Cell
MRD: Minimal Residual Disease
PBMC: Peripheral Blood Mononuclear Cell
RFs: Random Forests
SVMs: Support Vector Machines

## Supplementary Material

**Supplementary Figure 1:** Flowchart of Hypergate’s operating principle

**Supplementary Figure 2:** t-SNE representation of the ‘Samusik_01’ dataset color-coded by manual gating labels.

**Supplementary Figure 3:** Biplots of CD44, CD64 and F4/80 expression in the ‘Samusik_01’ dataset. Red events highlight C16, the t-SNE-defined macrophages.

**Supplementary Figure 4:** Biplots of CD64 versus F4/80 color-coded by intensity of either CD11b, SiglecF, B220, CD3 or Ly6C. These markers have been used by the authors to gate-out respectively monocytes, eosinophils, B cells and T cells.

**Supplementary Figure 5:** t-SNE representation of ALL samples at diagnosis (two first columns) or follow-up timepoints (three last columns) across five cases (rows). Blue denotes a malignant event manually identified and reviewed by an expert haematologist. Red denotes an event classified as malignant by Hypergate. The strategy learnt for column two is applied as is on columns three, four and five.

**Supplementary Video 1:** Animation of the optimization procedures applied to Non-Classical-Monocytes defined by manual clustering on the t-SNE biplot (showed as a dashed polygon). On the t-SNE biplot, events are plotted in either black if the gating procedure classifies them correctly, and in red otherwise. The right panels show the parameters selected in the final gating strategy, the red frame shows the cut-offs that are currently selected for these parameters, and a blue segment shows the next-chosen cut-off value. On the biplots, events are color-coded as black for True Positives, blue for False Negative, red for False Positive. True Negative are omitted. The bottom part shows the current value of the yield, purity and F1 score. F1 score steadily increases with each step, while purity increases during contractions, yield during expansions.

